# A South American mouse morbillivirus sheds light into a clade of rodent-borne morbilliviruses

**DOI:** 10.1101/2022.03.18.484959

**Authors:** Humberto J. Debat

## Abstract

Morbilliviruses are negative sense single stranded monosegmented RNA viruses in the family *Paramyxoviridae* (order *Mononegavirales*). Morbilliviruses infect diverse mammals including humans, dogs, cats, small ruminants, seals, and cetaceans, which serve as natural hosts. Here, I report the identification and characterization of novel viruses associated with South American long-haired and olive field mice. The divergent viruses dubbed Ratón oliváceo morbillivirus (RoMV) detected in renal samples from mice collected from Chile and Argentina are characterized by an unusually large genome including long intergenic regions and the presence of an accessory protein between the F and H genes redounding in a genome architecture consisting in 3’-N-P/V/C-M-F-hp-H-L-5’. Structural and functional annotation, genetic distance and evolutionary insights suggest that RoMV is a member of a novel species within genus *Morbillivirus* tentatively dubbed South American mouse morbillivirus. Phylogenetic analysis suggests that this mouse morbillivirus is closely related to, and clusters into a monophyletic group of novel rodent-borne morbilliviruses. This subclade of divergent viruses expands the host range, redefines the genomic organization and provides insights on the evolutionary history of genus *Morbillivirus*.

## Introduction

Genus *Morbillivirus* encompasses viruses in the family *Paramyxoviridae* (order *Mononegavirales)* affecting not only humans but also dogs, cats, small ruminants, seals, and cetaceans, that serve as natural hosts. Morbillivirus infection may cause diseases such as acute febrile respiratory tract infection in some animals and measles, a highly contagious infection disease in humans that triggers a severe immunosuppression, which generates over 140,000 vaccine-preventable deaths per year worldwide, mostly in sub-Saharan Africa (WHO, 2019). The etiological cause of measles is measles virus (MeV) from the species *Measles morbillivirus*, one of the seven species within the genus.

Morbilliviruses are enveloped with spherical virions harboring a single-stranded, negative-sense, monosegmented RNA genome. The envelope displays two glycoproteins with receptor attachment (haemagglutinin, H) and fusion functions (F). A matrix protein (M) is associated with the inner face of the envelope. The RNA genome is protected in a ribonucleoprotein core containing a nucleocapsid protein (N), a polymerase-associated co-factor protein (P) and a large protein (L), which is an RNA-directed RNA polymerase also involved in capping and cap methylation. Morbilliviruses have a genome ranging from 15,690–16,050 nt, with a canonical genomic architecture encoding eight proteins in the order 3′-N-P/V/C-M-F-H-L-5′ (Rima et al, 2019). The Morbillivirus genome organization includes a transcription unit with RNA editing encoding the phosphoprotein (P), a Zn2+-binding cysteine-rich protein (V) which is the edited mRNA form (one “G” added), and the overlapped non-structural protein (C) derived from leaky scanning.

Although morbilliviruses are hosted by several mammals, the apparently restricted host-range of members of each species is determined by receptors which define susceptibility of organisms to infection (Fukuhara, et al, 2019). For at least MeV, canine distemper virus (CDV) and rinderpest virus (RPV) the main receptors are signaling lymphocytic activation molecule family member 1 (CD150, SLAMF1).

Abundant novel viruses have been identified using metagenomic approaches, revealing the complexity of an expanding virosphere (Shi et al, 2018). RNA molecules of these viruses are often inadvertently co-purified with host RNAs and their sequences can be characterized from transcriptome datasets. In a consensus statement report, Simmonds et al (2017) indicate that viruses that are known only from metagenomic data can and have been incorporated as *bona fide* viruses into the official classification of the International Committee on Taxonomy of Viruses (ICTV). Thus, the analysis of transcriptome data represents an evolving source of novel virology insights, which allows the reliable identification of new viruses in hosts with no previous record of virus infections.

While there are numerous paramyxoviruses linked to rodents, intriguingly as of to date albeit their extended host range, there are no morbilliviruses where rodents unequivocally serve as natural main hosts. *Abrothrix hirta* and *Abrothrix olivacea* are sigmodontine rodents endemic to Chile and Argentina. Here, by analyzing public metagenomic data I report the identification and characterization of novel viruses associated to the South American long-haired and olive field mice representing a novel virus species from an emerging clade of rodent-borne morbilliviruses.

## Results and Discussion

In order to expand our knowledge on morbillivirus virus diversity I assessed an essential resource of public high-throughput sequencing RNA data available at NCBI: the Transcriptome Shotgun Assembly (TSA) Database available at https://www.ncbi.nlm.nih.gov/Traces/wgs/?view=TSA including 866 RNAseq datasets on diverse vertebrates. In tBLASTN searches (word size 6, expect threshold 10, scoring matrix BLOSUM62) against Vertebrata (taxid:7742) using as query the N protein of MeV (NP_056918.1) I retrieved a significant hit (E-value = 1e-142, 52.82% identity) corresponding to a 1,236 nt transcript from a TSA (GenBank: GCHM00000000.1) of a renal gene expression dataset of the South American long-haired mouse, *Abrothrix hirta* collected in natural populations (Valdez et al, 2015, BioProject PRJNA256304). *A. hirta* is a sigmodontine rodent widely distributed in southern South America occurring at both sides of the Andes, from the Chilean Region of Maule (ca. 35° S) to the Argentinean Tierra del Fuego (ca. 52° S). This specific library corresponded to half of kidney sample from an adult male (specimen PPA528) captured using Sherman live traps, three km west from Lago Pueyrredón in Río Chico department, Santa Cruz province, Argentina (47.42105 S, 71.958233 W) during field trips conducted during fall season (April 2011 and March 2012) (**Supplementary Figure 1**).

Further inspection by BLASTP searches (E-value < 1e-5) of the GCHM00000000.1 TSA library using as query MeV proteins retrieved four additional transcripts ranging from 462 to 1,922 nt which showed significant hits (E-value 3.43e-30 to 5.52e-134, identity 34.6% to 56%) to MeV encoded P, M and F proteins. The tentative virus contigs were curated by iterative mapping of the corresponding 76,503,290 library reads (NCBI-SRA: SRX663121) using Bowtie2 http://bowtie-bio.sourceforge.net/bowtie2/index.shtml with standard parameters which was also employed for mean coverage estimation and reads per million (RPMs) calculations. The transcripts extended, overlapped and polished by iterative cycles of mapping of raw reads were subsequently reassembled into a 9,941 nt long virus sequence including a continuum from a partial 3’ leader sequence followed by N-P/V/C-M-F-hp-H-and a few short partial sequences of L (1,720 nt long), with a mean coverage of 14.8x obtained with 1,779 virus derived 85×2 nt-long reads.

As this study collected a total of 16 adults of South American long-haired mice from Chile and Argentina I assessed by read mapping with Bowtie2 using as query the assembled virus sequence the 15 additional libraries, finding virus reads (82 and 62 total reads) in two. The samples where virus reads were detected (v+) matched to a female from the very same location (47.42105 S, 71.958233 W, specimen PPA357, SRA: SRX663109) and a male collected 65 km to the west in Aysén, Chile, (47.49671666 S, 72.80861666 W, specimen GD1454, SRA: SRX663075), respectively. While the number of reads was certainly low, inspection of the mapping results of virus reads from these additional libraries to the PPA528 consensus revealed some fixed SNPs among libraries (**Supplementary Figure 2**) suggesting: *i*) for the one hand that the reads were evidence of apparently three distinctive virus isolates. *ii*) On the other hand, ruling out that these few reads corresponded to spillover from the PPA528 sample or contamination artifact from index hoping during library processing.

In order to expand the survey of this virus to additional hosts I retrieved from NCBI all available RNAseq datasets of rodent subfamily *Sigmodontinae* (*Cricetidae*) including New World rats and mice, with at least 376 species. Of the 79 additional publicly available transcriptome datasets of mice including members of the *Sigmodon, Oligoryzomys* and *Abrothrix* genera (**Supplementary Table 1**) virus reads were detected in two libraries of *Abrothrix olivacea* (**Supplementary Figure 1**). The olive field mouse (*A. olivacea*) is the rodent that shows the broadest geographic distribution in southern South America. It ranges from the northernmost region of Chile (ca. 18° S), central-western Argentina (ca. 35° S) towards Patagonia, where it reaches the south of Tierra del Fuego (ca. 56° S). In elevation, it is found from sea level to up to 2,500 m of altitude (Giorello et al, 2014). Regarding viruses and abrotrichine rodents, to my knowledge, there are no reports oriented to the detection or characterizations of viruses linked to these mice. It is worth mentioning the detection of Andes hantavirus virus-reactive antibodies in a *A. olivacea* exemplars from southern Chile (Murrua, 1999) and that in experimental conditions the olive field mouse is susceptible to hantavirus infection (Padula et al, 2004).

The two specific v+ libraries corresponded to kidney samples from an adult male (specimen PPA444, SRA: SRX4099316) captured also in Río Chico department, Santa Cruz province, Argentina, but 265 km South East from the location where PPA528 and PPA357 were collected (49.42105 S, 69.958233 W). The other sample, also an adult male (specimen GD1411, SRA: SRX4099309) was captured 870 km to the North East, in Fundo San Martín, Región de Los Ríos, Chile (39.649233 S, 73.19255 W); both GD1411 and PPA444 were collected in the same study (BioProject PRJNA471316, Giorello et al, 2018). With iterative cycles of relaxed mapping (Bowtie2 parameters --very-sensitive-local) of SRX4099309 raw reads, extension and subsequent reassembly, a 9,948 nt long virus sequence from the GD1411 sample was obtained, including a continuum from a partial 3’ leader sequence to N-P/V/C-M-F-hp-H- and a few short partial sequences of L (2,736 nt long)), with a robust mean coverage of 128× obtained with 12,630 virus derived 101×2 nt-long reads. Notably, implementing the same pipeline to sample PPA444 employing the SRX4099316 library a coding complete virus sequence with the genome architecture 3’-N-P/V/C-M-F-hp-H-L-5’ was assembled corresponding to 16,568 nt supported by a 17.7× mean coverage from 2,897 virus derived 101×2 nt-long reads.

A rapid comparison based on sequence alignments of the three consensus virus sequences (**Supplementary Figure 3**) indicated that while divergent, the % identity of predicted proteins ranged between 88.2-99.8% suggesting that the sequences corresponded to three distinctive strains of the same virus which I tentatively dubbed Ratón oliváceo morbillivirus (RoMV). More in detail, the viruses assembled from the PPA444 and PPA528 samples are highly similar with their ORFs and predicted proteins sharing over a 99% sequence identity. In contrast, the GD1411 virus is clearly more divergent sharing a lower 85-89% nt identity and 88-98% aa identity of the predicted proteins, being the P and H proteins the more distinctive and N and M the more similar, indicating the significant preeminence of synonymous mutations on the GD1411 virus. Perhaps is worth emphasizing that the GD1411 mouse was collected on the other side of the Andes mountain range, over 850 km and more than 1,100 km North East of the places where PPA528 and PPA444 were captured, signifying that geographical isolation could provide some clues in the evolutionary history of these viruses. The significant diversity revealed by these three mouse viruses could indicate a long lasting virus-host relationship between RoMV and abrotrichine rodents. In turn, the consensus assembly from sample PPA444 that comprised a complete coding, (near) complete genome was used as reference for structural and functional annotation, genomic comparison and evolutionary insights of RoMV.

To characterize thoroughly the RoMV sequence, virus annotation was implemented as reported elsewhere (Debat, 2017; Debat & Bejerman, 2022). In brief, virus open reading frames (ORFs) were predicted with ORFfinder (https://www.ncbi.nlm.nih.gov/orffinder/) domains presence and architecture of predicted proteins was determined by the NCBI Conserved domain database v3.16 (https://www.ncbi.nlm.nih.gov/Structure/cdd/wrpsb.cgi) and HHPred/HHBlits available at https://toolkit.tuebingen.mpg.de/#/tools/. Secondary protein structure was predicted with Garnier http://emboss.sourceforge.net/apps/release/6.6/emboss/apps/garnier.html, signal peptides were assessed with SingnalP v5 https://services.healthtech.dtu.dk/service.php?SignalP-5.0 and prediction of transmembrane topology with Phobius https://phobius.sbc.su.se/.

The tentatively named Ratón oliváceo morbillivirus genome organization is characterized by a □16,658 nt long negative-sense single-stranded RNA containing six main ORFs in the anti-genome, positive-sense orientation. In addition the second ORF includes a transcription unit with RNA editing and an overlapping ORF (P/V/C) and between the F and H genes there is an additional accessory ORF. In sum the genomic architecture of RoMV is 3’-N-P/V/C-M-F-hp-H-L-5’ (**Figure 1.A**). As expected for paramyxoviruses the genes are separated by intergenic gene junctions regions, composed of the polyadenylation signal of the preceding gene, a short intergenic region, and the transcriptional start of the following gene (Rima et al, 2019). The detected consensus gene junction region of RoMV is consistent with morbilliviruses which have a conserved intergenic motif (CUU) between the gene-end and gene-start of adjacent genes following the structure “AAAA-CUU-AGG” (**Table 1**).

**Figure 1.**
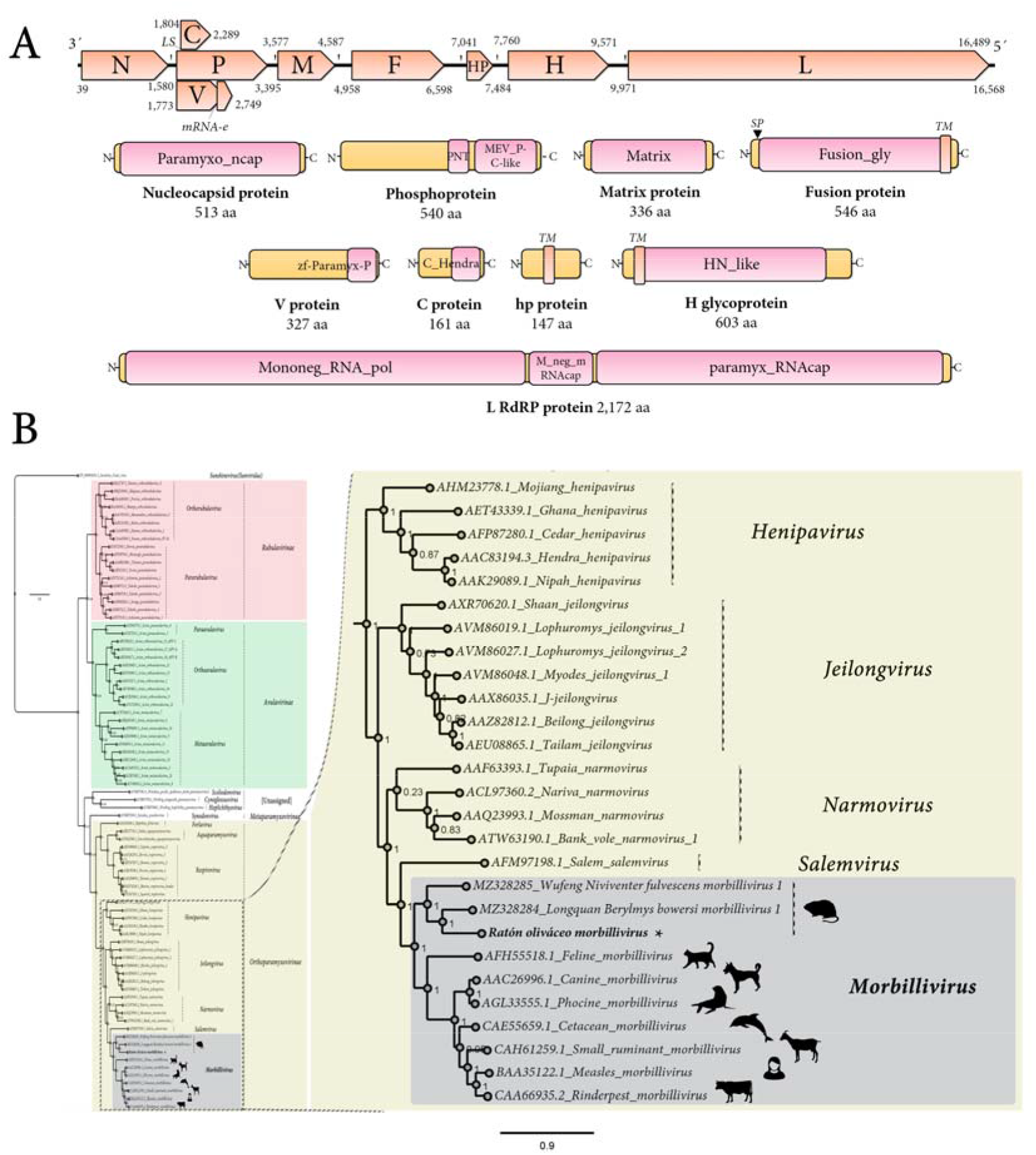
Genomic architecture and phylogenetic insights of Ratón oliváceo morbillivirus. (**A**) Genome graphs depicting architecture and predicted gene products of RoMV. The predicted coding sequences are shown in orange arrow rectangles, start and end coordinates are indicated. Gene products are depicted in curved yellow rectangles and size in aa and name is indicated below. Predicted domains or HHPred best-hit regions are shown in curved pink rectangles. Abbreviations: N, nucleoprotein CDS; P, phosphoprotein CDS; M, Matrix protein CDS; F, Fusion protein CDS; HP, hypothetical protein CDS, H, Hemagglutinin glycoprotein CDS; L, RNA dependent RNA polymerase CDS; TM, trans-membrane domain; SP, signal peptide; LS, leaky scanning; mRNA-e, mRNA editing. Domain abbreviations are described in the main text. (**B**) Maximum likelihood phylogenetic tree based on amino acid alignments of the L polymerase of RoMV and members of accepted species within family *Paramyxoviridae*. The right panel represents an inset of the complete phylogenetic tree which is available as **Supplementary Figure 9**. The tree is rooted at Sunshine coast virus (family *Sunviridae*). The scale bar indicates the number of substitutions per site. Node labels indicate FastTree support values. Silhouettes represent natural hosts. Genbank accession numbers of the proteins used are depicted before each virus species name.

BLASTP searches of predicted products (**Table 2**) tentatively identified these ORFs as potentially encoding: a nucleocapsid protein (N; 513 aa), phosphoprotein (P; 540 aa), V non-structural protein (V; 325 aa), C non-structural protein (C; 161 aa), a matrix protein (M; 336 aa), a fusion protein (F; 546 aa), a small hypothetical protein (hp; 147 aa), a Hemagglutinin glycoprotein (H; 603 aa), and an RNA dependent RNA polymerase (L; 2,172 aa). Importantly, all best hits based on highest sequence identity scores, which ranged between 25.1% (H) to 63.7% (M), were morbilliviruses, more specifically MeV, Longquan Berylmys bowersi morbillivirus 1 (LBbMV) and Wufeng Niviventer fulvescens morbillivirus 1 (WNfMV). LBbMV and WNfMV correspond to recently released virus sequences, which are unpublished yet and have been annotated as unclassified morbilliviruses. The metadata of their GenBank accessions indicate that LBbMV (MZ328284) was identified in Bower’s white-toothed rat (*Berylmys bowersi*), a rodent from family *Muridae* which is native to Southeast Asia. WNfMV (MZ328285) was detected in chestnut white-bellied rat (*Niviventer fulvescens*) another rodent from family *Muridae*.

Structural and functional annotation indicates that the 513 aa RoMV-N protein harbors a Paramyxovirus nucleocapsid protein domain (Paramyxo_ncap, E-value = 0, coordinates 1-512) which is involved in tightly encapsidating the viral RNA and interacting with several other viral encoded proteins, all of which are involved in controlling replication. RoMV-N presents the conserved MA(S,T)L motif of morbilliviruses, and appears to share the three key conserved motifs in paramyxoviruses and nuclear export signals and NLS (**Supplementary Figure 4**). The 540 aa P protein, which plays a crucial role by positioning L onto the N/RNA template through an interaction with the C-terminal domain of N is a co-factor of the RdRP, includes a Paramyxovirus structural protein V/P N-terminus domain (Paramyxo_PNT, pfam13825, E-value = 3.67e-03, coordinates 274-347), and a Paramyxovirus P/V phosphoprotein C-terminal domain (Paramyx_P_V_C, pfam03210, E-value = 1.13e-16, coordinates 372-536) (**Figure 1.A**). Most of its 540 amino acids, as expected, appear to be in a natively disordered state, and the C-terminal conserved residues are putatively folded into a three-helical bundle that binds to the C-terminal tail of N and has an oligomerization domain that forms a long tetrameric coiled-coil that is stabilized at its N terminus by a helical bundle linking protomers. 3d modelling using the Swiss-Model platform available at https://swissmodel.expasy.org/ with standard parameters using as best fit template the 3zdo.1.B MeV phosphoprotein (Communie et al, 2013) showed that RoMV-P forms a tetrameric coiled coil with the similar length (63 aa) and conserved structure but less highly packed as the measles virus P protein (**Supplementary Figure 5**). The 327 aa cysteine-rich non-structural V protein generated by mRNA editing by incorporating an additional “G” at coordinate 2,512 of what encodes the P mRNA has a Zinc-binding domain of *Paramyxoviridae* V protein at its C-terminal region (zf-Paramyx-P, E-value = 2.4e-15, coordinates 280-323). V is generated by an A-rich context where the RNA transcriptase ‘stutters’ on the template at the editing motif which is “AAAAAGGG” in RoMV. This stuttering results in the insertion of one pseudo-templated G shifting the reading frame to access the alternative ORF V (Rima et al, 2019). The 161 aa C protein, which is generated by leaky scanning of the P mRNA which results in the translation of an overlapped ORF 31 nt downstream the AUG of P, presents a C protein from hendra and measles viruses domain (C_Hendra, pfam16821, E-value = 8.10e-05, coordinates 1-146). C protein has been involved in host defense interaction, for instance MeV C is implicated in modulation of interferon signaling but also in pathogenicity and virulence as is the case for CDV C (Siering et al, 2020). The non-glycosylated membrane or matrix protein (M) is 336 aa long and has a viral matrix protein domain (Matrix, pfam00661, E-value = 6.34e-118, coordinates 6-326). M appears to be the most conserved protein of RoMV, sharing 63.6% aa identity to that of LBbMV. The 546 aa F protein presents as expected a signal peptide at its N-terminal region and a transmembrane domain at its C-end (**Figure 1.A**). F functional annotation pinpointed a typical Fusion glycoprotein domain (Fusion_gly, pfam00523, E-value = 1.04e-128, coordinates 23-478). Unexpectedly, a small 147 aa protein (hp) was found between the F-H intergenic region showing no homology to any protein, nor domains. No similarities were found to motifs/domains/peptides/proteins in any database to hp when Psi-blast, HHblits, HHPred or HMMER searches were implemented (see below for more details). The 603 aa surface H glycoprotein showed a typical Haemagglutinin-neuraminidase of paramyxoviridae domain (HN_like, cd15464, E-value = 3.09e-20, coordinates 207-579) and a N-terminal transmembrane domain. The H protein is the most divergent encoded main protein RoMV showing only a 16-22% aa best identity with the H of LBbMV and WNfMV.

As CD150 is the tentative main receptor of morbilliviruses I used the primary data from *A. olivacea* to reconstruct the protein using as query the signaling lymphocytic activation molecule family member 1 coding sequence from the available hispid cotton rat CD150 (*Sigmodon hispidus*, JX424845), eventually generating a complete mRNA 1,278 nt long encoding a *A. olivacea* 340 aa protein showing 81.5% aa identity to that of hispid cotton rat. In order to try to glimpse the RBP-CD150 interactions that could be involved in determining host tropism, I compared the amino acid sequences at the putative contact surfaces of morbillivirus RBPs and their cognate CD150 receptors based on the predictions of Ikegame et al (2021). Alignment of putative key regions in some morbillivirus H proteins implicated in CD150 interactions showed virus-specific changes, with some residues highly conserved and others significantly variable, that may suggest adaptation of morbillivirus H to the putative CD150 receptors of their cognate host (**Supplementary Figure 6**). Modelling and experimental assessment of these *in silico* predictions could provide some lights in the specific role of *A. olivacea* CD150 in host interaction and range of RoMV. Finally, the 2,172 aa long L protein presents a Mononegavirales RNA dependent RNA polymerase domain (Mononeg_RNA_pol, pfam00946, E-value = O, coordinates 16-1107) followed by a *Paramyxoviridae* family mRNA capping enzyme region (paramyx_RNAcap, TIGR04198, E-value = 2.96e-176, coordinates 1224-2172) including a mRNA (guanine-7-)methyltransferase (G-7-MTase) (G-7-MTase, pfam12803, E-value = 1.74e-87, coordinates 1483-1793) which catalyzes cap methylation.

The genome of RoMV presents some peculiarities that distinguish it from other assigned morbilliviruses. For instance, the (nearly) complete sequence with 16,658 nt represents to date the lengthiest morbillivirus reported (**Supplementary Figure 7**), being as is at least 518 nt longer than Feline morbillivirus (FeMV) which is characterized for a long M-F intergenic region and 5’ trailer sequence. The longer nature of RoMV is not explained by its coding regions, which are of typical size, but by the longest intergenic regions described yet within the genus (**Supplementary Table 2**). RoMV presents the lengthiest N-P, P-M and F-H intergenic regions reported yet. In the latter, the presence of an accessory putative ORF encoding a 147 aa hypothetical protein with a trans membrane domain is not a hallmark of morbilliviruses. It is worth noting that the rodent putative morbillivirus WNfMV shares in the same genomic context between F-H an ORF encoding a small 74 aa protein also with a transmembrane domain. While it is tempting to consider that these accessory proteins may have some role with rodent host interaction, the absence of this ORF in LBbMV hampers any conclusion, its function remains elusive, and its presence is not a distinguishing feature of this subclade. A short integral membrane protein (SH) and/or transmembrane protein (tM) located between F-H is not exceptional in paramyxoviruses and can be found for instance in some members of subfamily *Orthoparamyxovirinae* such as rodent viruses from genus *Jeilonvirus* where it is thought to be involved in cell-to-cell fusion (Vanmechelen et al, 2018). Besides genomic location, relative size and presence of a transmembrane signal, there is no apparent identity or reminiscence of homology between these jeilonvirus proteins and the one from RoMV and WNfMV, thus I decided to dub it as hypothetical protein (hp) instead of SH or tM to avoid confusions (**Supplementary Figure 8**). RoMV and LBbMV include also a significantly long H-L intergenic region of about three times the typical size in morbilliviruses mainly derived from an unusually long AU rich (65-70%) H mRNA 3’UTR.

Phylogenetic insights based on the predicted replicase of RoMV were employed to assess the putative evolutionary placement of this virus. To this end, the L protein aa alignment of recognized members of the family *Paramyxoviridae* provided as a resource of ICTV available at https://talk.ictvonline.org/ictv-reports/ictv_online_report/negative-sense-rna-viruses/w/paramyxoviridae/1197/resources-paramyxoviridae was retrieved and a consensus alignment was generated using ClustalW with gap generation penalties of five and extension penalties of one in both multi- and pairwise alignments including RoMV, WNfMV and LBbMV. Additionally, predicted proteins N, P, M, F and H alignments were generated by MAFTT 7.490 https://mafft.cbrc.jp/alignment/software/ (BLOSUM62 scoring matrix) using as best-fit algorithm E-INS-i (M and P) or G-INS-i (N, F and H). The aligned proteins were used as input for FastTree 2.1.11 (http://www.microbesonline.org/fasttree/) maximum likelihood phylogenetic trees (best-fit model = JTT-Jones-Taylor-Thorton with single rate of evolution for each site = CAT) computing local support values with the Shimodaira-Hasegawa test (SH) and 1,000 tree resamples. The obtained paramyxovirus L tree clearly shows that RoMV clusters together with other viruses within the genus *Morbillivirus* (**Figure 1.B**; **Supplementary Figure 9**). In addition, RoMV appears to have a close evolutionary relationship with WNfMV and LBbMV, branching together forming a clade of rodent morbilliviruses. To further confirm these findings N, P, M, F and H phylogenetic trees were generated using proteins from viruses of genus *Morbillivirus*, RoMV, WNfMV, LBbMV and the respective proteins of Tupaia narmovirus and Nariva narmovirus (genus *Narmovirus*). In all cases, unequivocally, RoMV clustered with morbillivirus forming a sub-clade with the rodent WNfMV and LBbMV (**Supplementary Figure 10**). It is essential to highlight that during the preparation of this work a putative morbillivirus was described linked to wood mouse (*Apodemus sylvaticus*) a murid rodent native to Europe (Vanmechelen et al, 2022). While the genome sequence of this “Apodemus morbillivirus” detected in a wood mouse cadaver that had been killed by cats or vehicles collected in Belgium is yet publicly unavailable, its reported phylogenetic clustering with both WNfMV and LBbMV is consistent with a the novel sister subclade of rodent-borne morbilliviruses described here.

The ICTV species demarcation criteria of morbilliviruses is based on distance in the phylogenic tree of complete L protein based on tree topology and branch length between the nearest node and the tip of the branch. This length is defined as 0.03 in the trees generated in the ICTV paramyxovirus resource and used as input for the L tree reported here. As the branch length from the node separating RoMV/LBbMV in substitutions per site of the obtained consensus tree is well above this threshold, RoMV appears to correspond to a new virus, a putative member of a novel species within genus *Morbillivirus* which I tentatively name ‘South American mouse morbillivirus’.

## Conclusions

In summary, the scrutiny of public NCBI-SRA transcriptome data constitutes a reliable source for the detection of novel viruses. By implementing a simple pipeline of RNA sequence analysis, I report the identification and molecular characterization of a novel rodent virus. The analyses revealed that these viruses are tentative members of a novel species of the genus *Morbillivirus*. The mouse morbillivirus reported here in parallel with additional recent reports and posted sequences sheds light on an emerging novel subclade of rodent morbilliviruses. More importantly, these novel viruses expand the genome architecture and host range of morbilliviruses providing new clues on the understanding of the evolutionary history of this genus of viruses. Future studies should assess additional rodent viruses, the potential prevalence of RoMV or related viruses in *Sigmodontinae* rodents, unravel whether the infection of these novel viruses is associated to specific symptoms, and explore the zoonotic and pathogenic potential of these viruses.

## Data availability

The virus sequences assembled in this study corresponding to Ratón oliváceo morbillivirus (RoMV strains PPA444, GD1411 and PPA528) have been submitted to NCBI-GenBank TPA and are available as **Supplementary Material 1** of this article. The primary data underlying these assemblies is publicly available on NCBI-BioProjects PRJNA471316 and PRJNA256304 and SRA experiments SRX4099316, SRX4099309 and SRX663121.

## Acknowledgments

I would like to express a sincere gratitude to the generators of the underlying data used for this work: Dr. Enrique P. Lessa and Dr. Guillermo D’Elía. By following open access practices and supporting accessible raw sequence data in public repositories available to the research community, they have promoted the generation of new knowledge and ideas.

## Compliance with ethical standards

### Ethical Approval

This article does not contain any studies with human participants or animals performed by any of the authors.

### Funding

This research did not receive any specific grant from funding agencies in the public, commercial, or not-for-profit sectors.

---

**Supplementary Figure 1.**
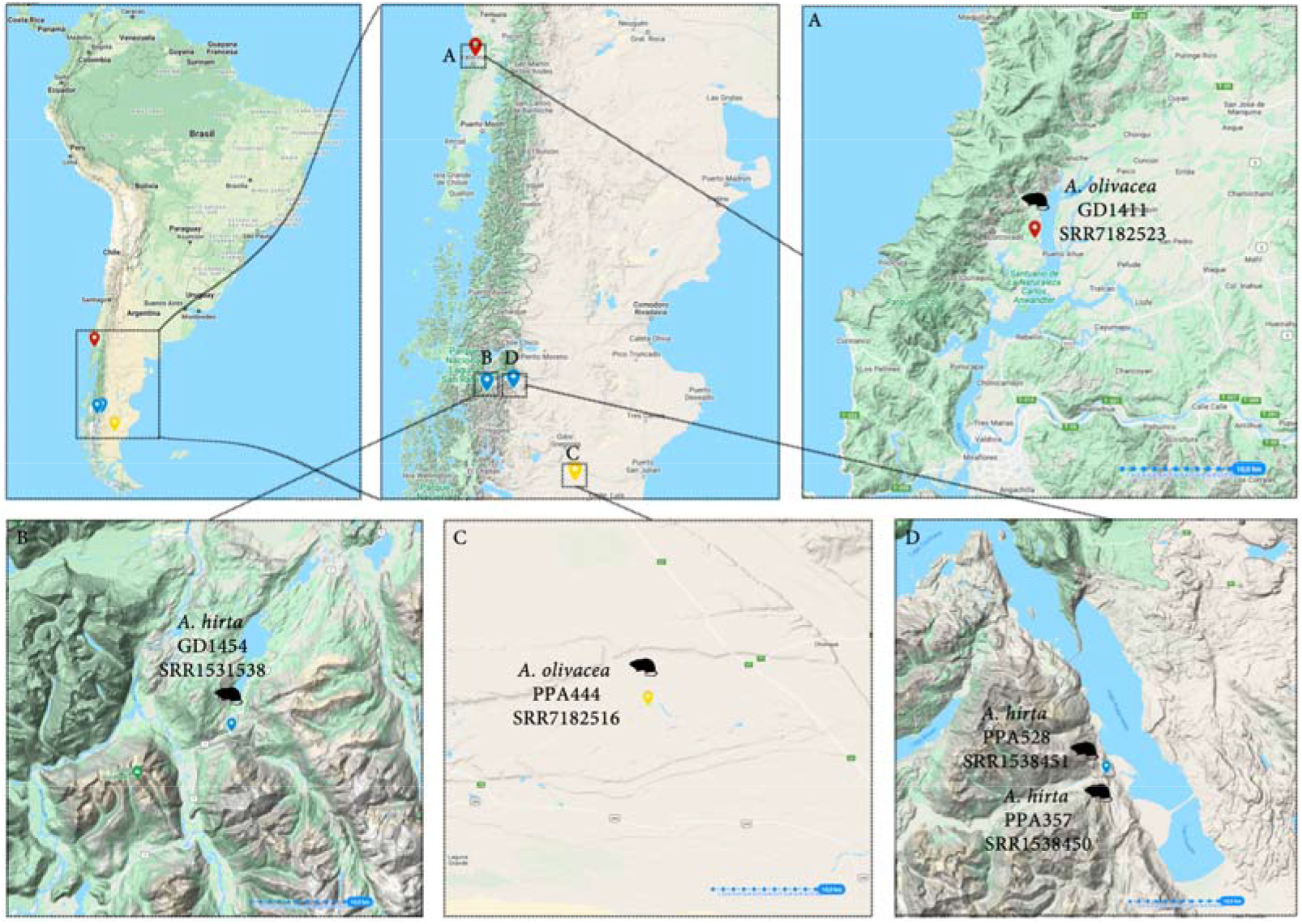
South American map depicting the location (pinned) of the captured mice where RoMV was detected. Inset of each specific region are expanded and ruler (km) is represented by a blue discontinued line at the right bottom.

**Supplementary Figure 2.**
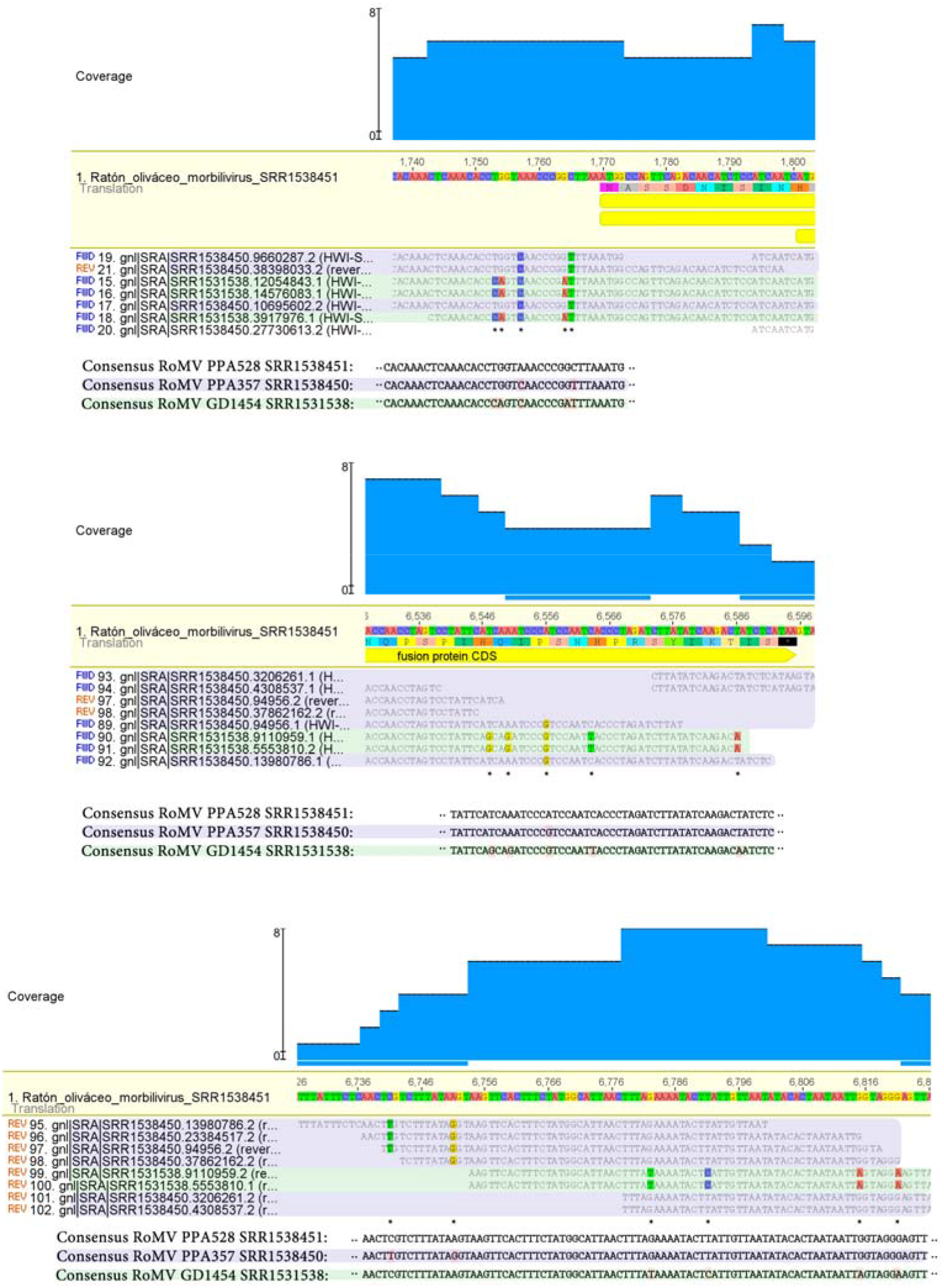
Mapping of RoMV PPA357 and GD1454 virus reads against the PPA528 consensus assembly. Three representative regions are depicted showing apparent fixed SNPs which differentiate the emerging consensus sequence of each sample. Reads from samples PPA357 and GD1454 are colored in light blue and green respectively. Asterisks indicate polymorphic sites.

**Supplementary Figure 3.**
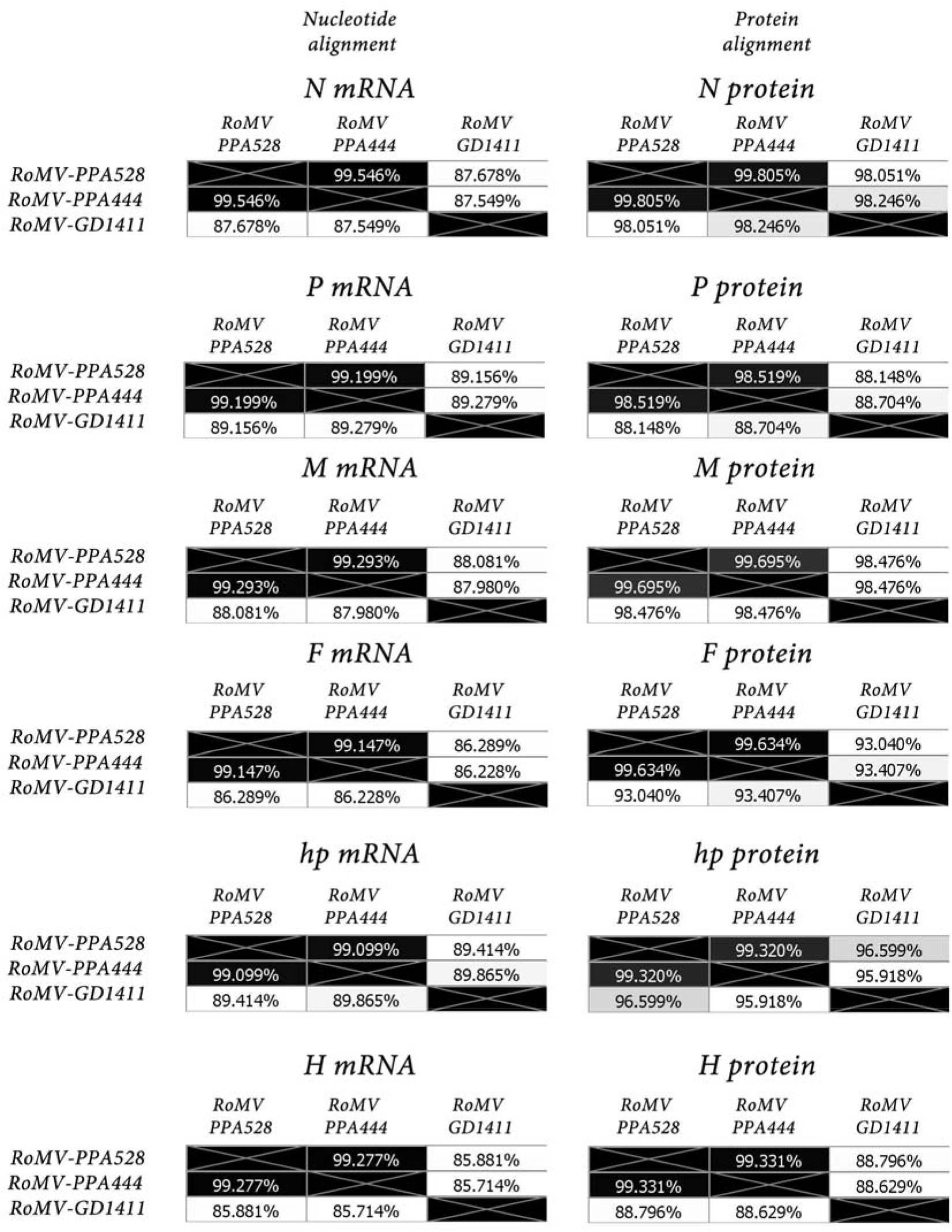
Genetic distance comparison of RoMV strains. Heat maps of % identity of nucleotide alignment of each mRNA (left panels) or protein alignments (right panels). Shading from white to black indicate lowest to highest sequence identity. L mRNA and protein is excluded given that the assemblies of RoMV PPA444 and GD1411 include only partial and dispersed encoding region of the RdRP.

**Supplementary Figure 4.**
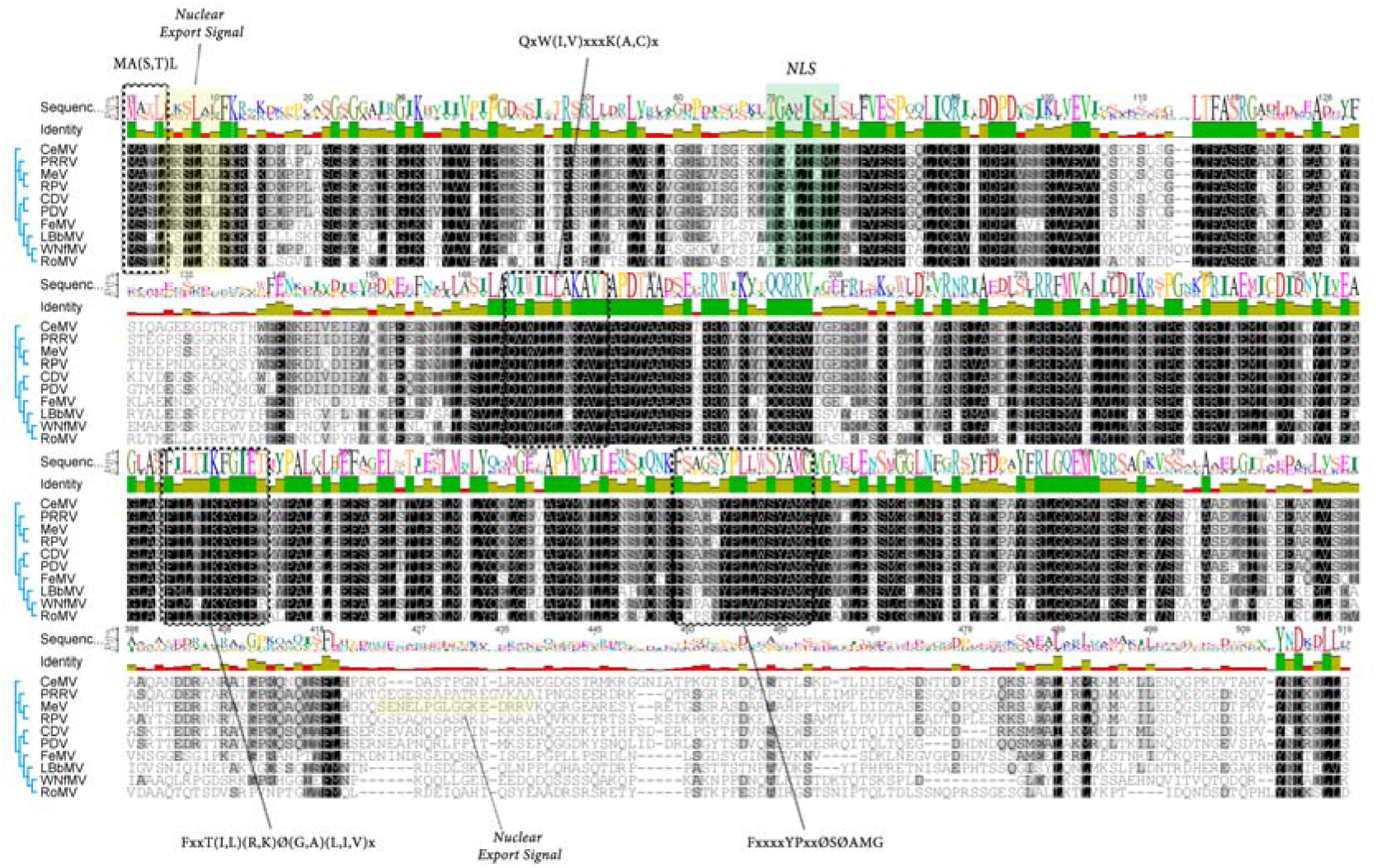
Multiple alignments of N proteins of RoMV and other morbilliviruses and recently described putative rodent morbilliviruses. The conserved MA(S,T)L motif in morbilliviruses and the three conserved motifs in paramyxoviruses are marked in open boxes with discontinued lines and consensus sequences are indicated above the alignment. “Ø” represents an aromatic aa and “x” represents any aa. The nuclear export signals are highlighted in yellow and the NLS highlighted in green.

**Supplementary Figure 5.**
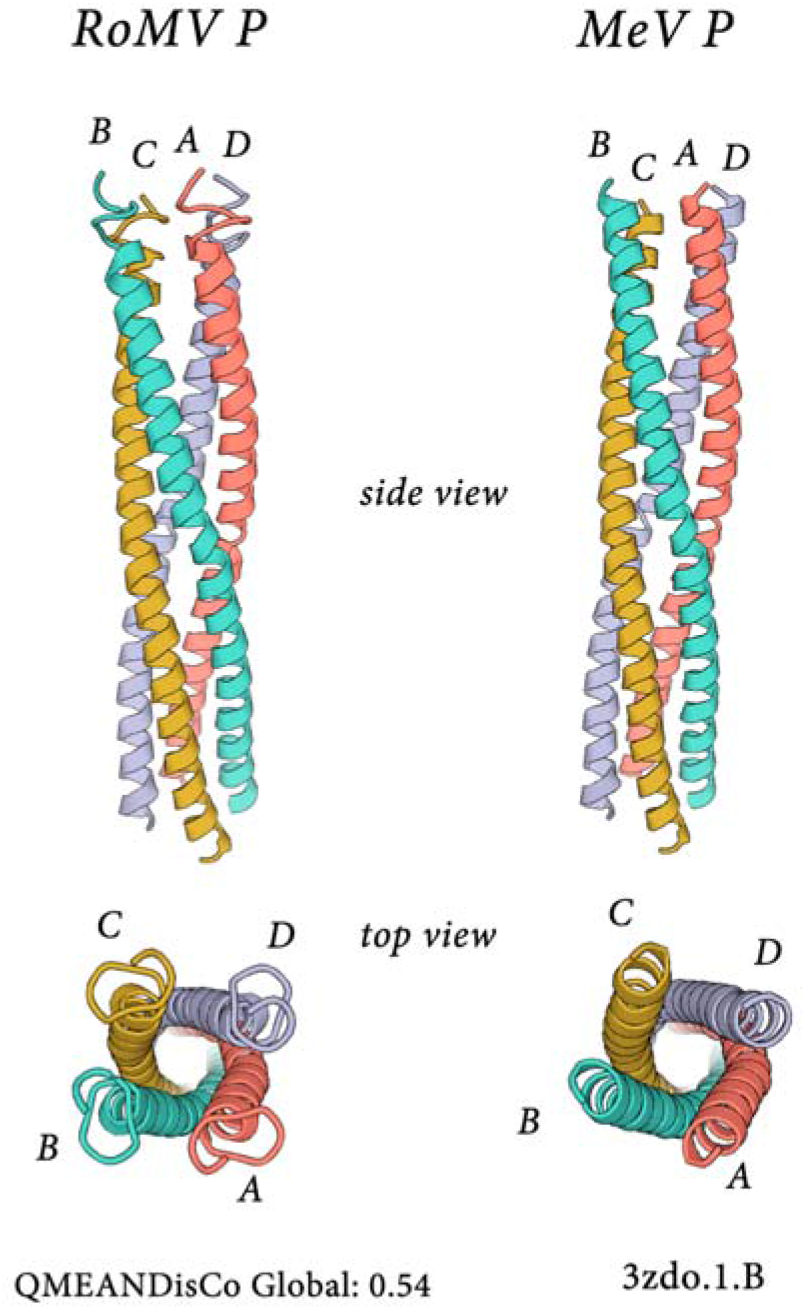
Side and top views of cartoon representations of the P oligomerization domains of RoMV and measles virus (MeV) generated by the SWISS-MODEL platform using as best fit template the 3zdo.1.B phosphoprotein. Letters A-D represent the putative monomers.

**Supplementary Figure 6.**
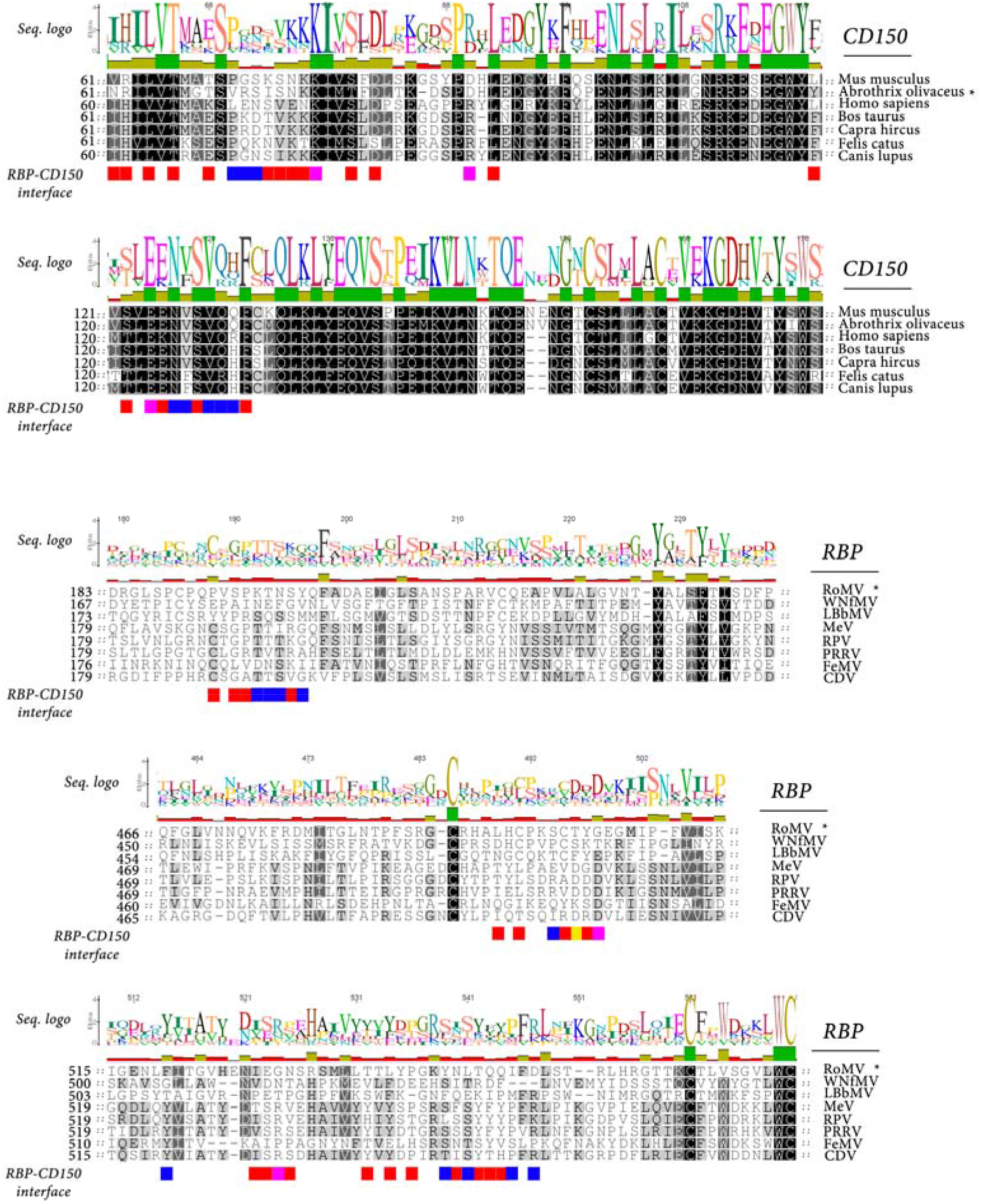
(Upper panel) AA sequence alignment of CD150 from some morbillivirus hosts and *A. olivacea*. CD150 from the 5 mammals that are the natural hosts, Mus musculus and olive field mouse aligned by clustalw. Alignments around the RNA Binding Domain (RBP) contact residues (aa60-179) are shown. Highly and moderately conserved residues in these regions are shaded in black and gray, respectively. Occluded residues at the RBP-CD150 interface (PDB: 3ALZ) are indicated by red color and other shading below the alignment. Hydrogen bonds (deep blue), salt bridges (yellow), hydrogen bonds + salt bridge (violet) in these occluded regions are indicated. (lower panel) AA sequence alignment of morbillivirus H proteins. Alignments around the CD150 contact residues are shown separately.

**Supplementary Figure 7.**
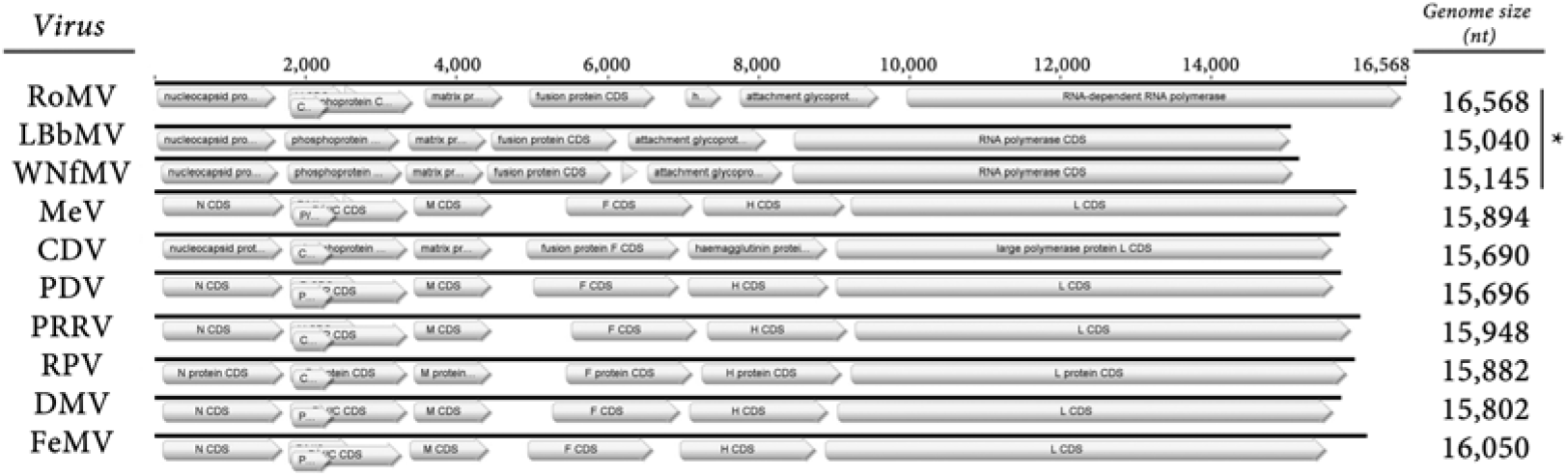
Genome graphs drawn at scale depicting accepted morbilliviruses, WNfMV, LBbMV and RoMV. ORFs are depicted as grey arrowed rectangles; annotations abbreviations are clarified in the main text. Virus name abbreviations are included in Supplementary Table 2.

**Supplementary Figure 8.**
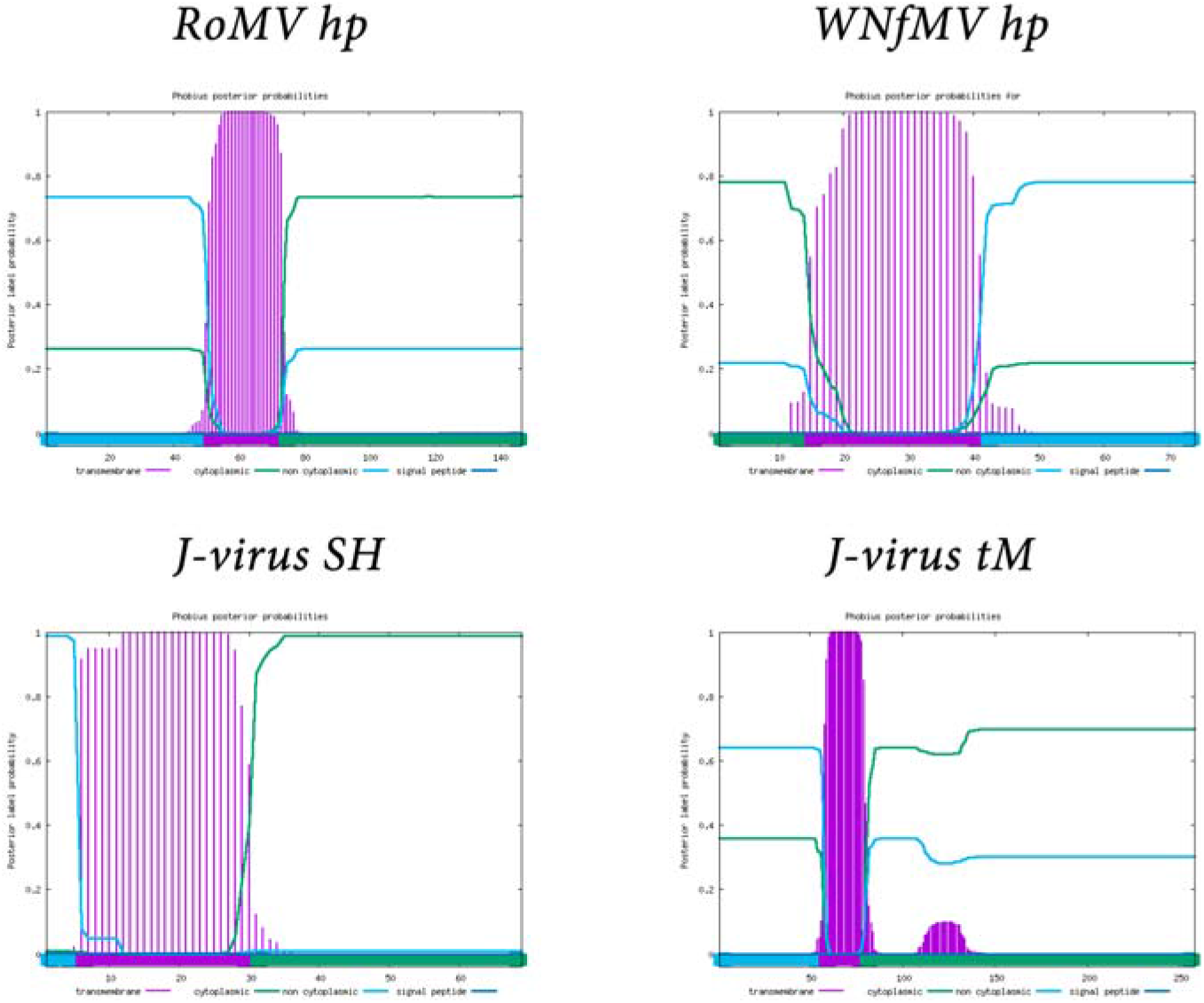
Diagrams representing **t**ransmembrane topology of hypothetical protein of RoMV and WNfMV and of J-virus (jeilonvirus) SH and tM as predicted by the Phobius tool.

**Supplementary Figure 9.**
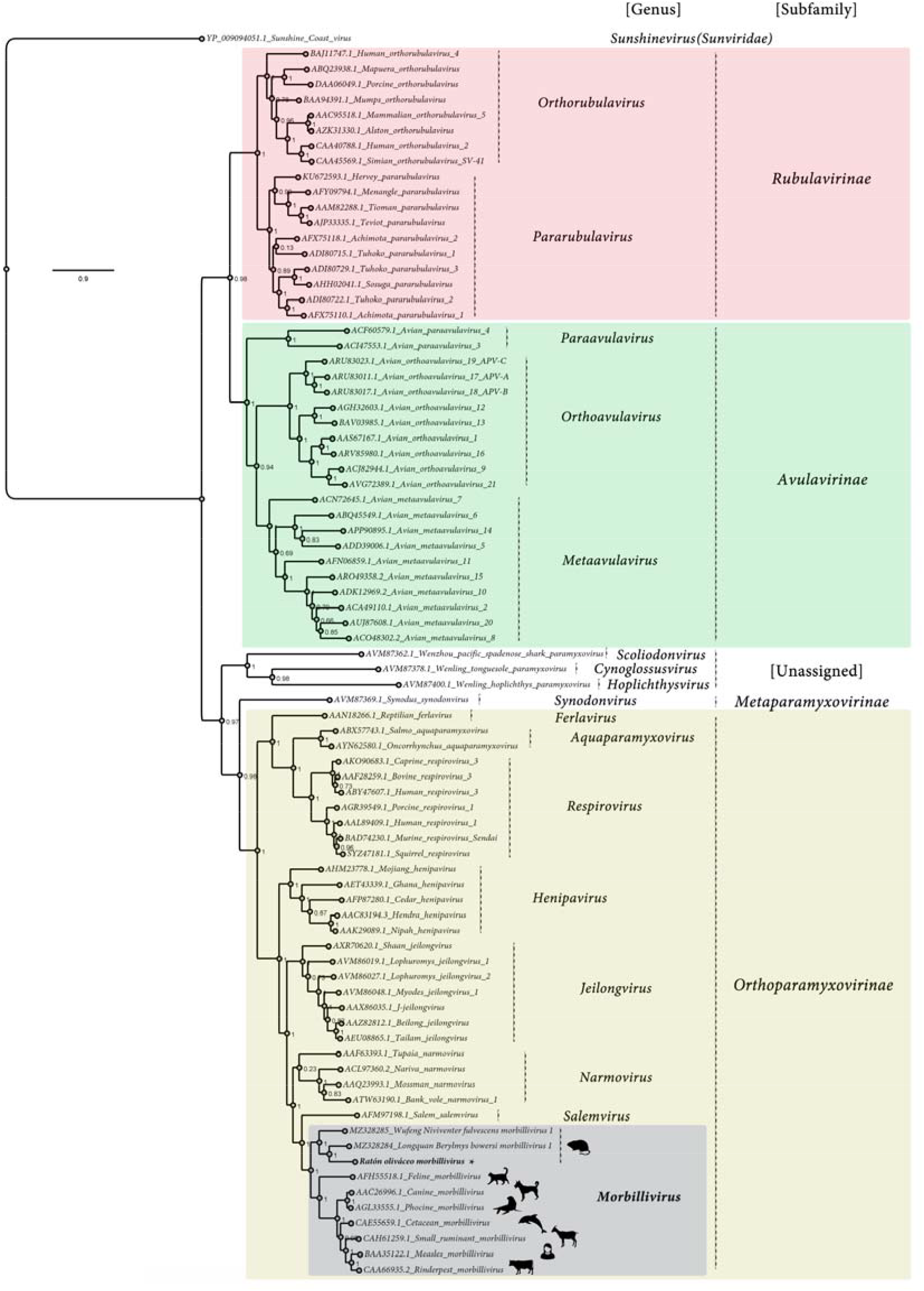
Maximum likelihood phylogenetic tree based on amino acid alignments of the L polymerase of RoMV and members of accepted species within family *Paramyxoviridae*. The tree is rooted at Sunshine coast virus (family *Sunviridae*). The scale bar indicates the number of substitutions per site. Node labels indicate FastTree support values. Silhouettes represent natural hosts. Genbank accession numbers of the proteins used are depicted before each virus species name.

**Supplementary Figure 10.**
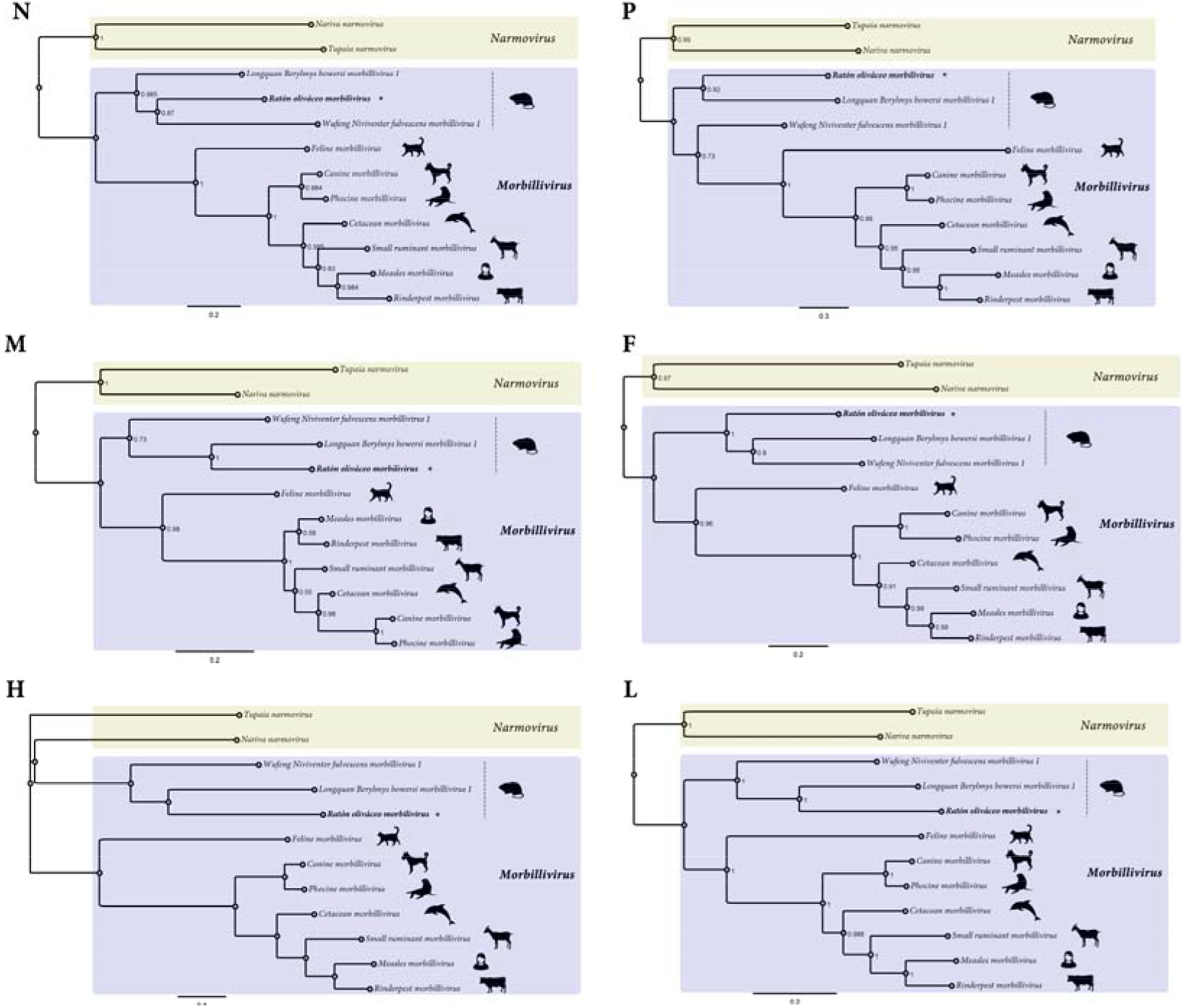
Maximum likelihood phylogenetic tree based on amino acid alignments of the N, P, M, F, H and L proteins of RoMV and members of accepted species within genus *Morbillivirus*, WNfMV, LBbMV (blue rectangle) and of Tupaia narmovirus and Nariva narmovirus (genus *Narmovirus*) (yellow rectangles). The scale bar indicates the number of substitutions per site. Node labels indicate FastTree support values. Silhouettes represent natural hosts. Letters at the upper left corner indicate the protein used for each tree.

