## Supplementary material for "A South American mouse morbillivirus sheds light into a clade of rodent-borne morbilliviruses": Table 2

| **ORF** | **Gene** | **Putative function** | **Lenght (aa)** | **MW (kD)** | **IEP^a^** | **BLASTP e-value** | **BLASTP Ide(%)** | **BLASTP Score** | **Best Hit abbr** |
| --- | --- | --- | --- | --- | --- | --- | --- | --- | --- |
| 1 | N | Nucleocapsid protein | 513 | 57.2 | 4.9 | 0.0 | **54.8** | 542 | LBbMV^b^ |
| 2 | P | Phosphoprotein | 540 | 60.2 | 5.1 | 3e-58 | **30.6** | 212 | LBbMV^b^ |
| 3 | V | V protein | 327 | 36.1 | 5.7 | 1e-15 | **35.5** | 87.4 | MeV^c^ |
| 4 | C | C protein | 161 | 18.8 | 11.1 | 0.037 | **37.0** | 44.7 | BvV1^d€^ |
| 5 | M | Matrix protein | 336 | 37.3 | 9.1 | 7e-154 | **63.7** | 445 | LBbMV^b^ |
| 6 | F | Fusion protein | 546 | 59.9 | 7.4 | 6e-178 | **53.0** | 523 | LBbMV^b^ |
| 7 | HP | Hypothetical protein | 147 | 16.8 | 9.6 | nh^e^ | nh^e^ | nh^e^ | nh^e^ |
| 8 | H | Glycoprotein | 603 | 66.7 | 7.7 | 5e-46 | **25.1** | 182 | LBbMV^b^ |
| 9 | L | RdRNA Polymerase | 2,172 | 248.7 | 7.1 | 0.0 | **62.8** | 2,915 | LBbMV^b^ |

**Table 2**. Summary of RoMV genome encoded proteins and best blasp hits.

^a^IEP, isoelectric point pH. ^b^LBbMV, Longquan Berylmys bowersi morbillivirus 1. ^c^MeV, Measles morbillivirus. ^d^BvV1, bank vole virus 1. ^e^nh, no hit. ^€^while the best obtained blastp hit of RoMV C protein retrieved BvV1, RoMV C closest identity corresponds to the unannotated overlapped gen product of LBbMV with shares with RoMV C a 37.4% aa identity.
