## Supplementary material for "A South American mouse morbillivirus sheds light into a clade of rodent-borne morbilliviruses": Table 1

**Table 1**. RoMV genome predicted intergenic regions.

|  | **Poly-adenylation** | **Intergenic spacer** | **Transcription start** |  |
| --- | --- | --- | --- | --- |
| 3′ leader | na | na | AGGTGCCAGC | N gene |
| N gene | ATTAAGAAAAA | CTT | AGGACCAAAG | P gene |
| P gene | ACTAAAGAAAA | CTT | AGGATTTAAA | M gene |
| M gene | ATTCAATAAAA | CTC | AGAGAATCTA | F gene |
| F gene | ATTAAGAAAAA | CTT | AGGAGGTAAA | HP gene |
| HP gene | ACTAAAGAAAA | CTT | AGGGTTAATG | G gene |
| G gene | TAAAGAAAACA | CTT | AGGAATAACG | L gene |
| L gene | ATTAAGAAAAA | CTT | na | 5′ leader |
| Consensus 85% | AYTAARnAAAA | CTT | AGGRnnWAnR |  |
| Consensus 100% | WHWMRRnAAMA | CTY | AGRDnnHMnV |  |
| Consensus MeV | VHHWHDnAAAA | CKT | AGGRnRMARG |  |
